# Sec17 (α-SNAP) and an SM-tethering complex control the outcome of SNARE zippering *in vitro* and *in vivo*

**DOI:** 10.1101/123133

**Authors:** Matthew L. Schwartz, Daniel P. Nickerson, Braden T. Lobingier, Cortney G. Angers, Michael Zick, Alexey J. Merz

## Abstract

Zippering of SNARE complexes spanning docked membranes is essential for most intracellular fusion events. Here we explore how SNARE regulators operate on discrete zippering states. The formation of a metastable *trans*-complex, catalyzed by HOPS and its SM subunit Vps33, is followed by subsequent zippering transitions that increase the probability of fusion. Operating independently of Sec18 catalysis, Sec17 either inhibits or stimulates SNARE-mediated fusion. If HOPS or Vps33 are absent, Sec17 inhibits fusion at an early stage. Thus, HOPS and Vps33 accelerate SNARE zippering, particularly in the presence of otherwise inhibitory Sec17. Once SNAREs are partially-zipped, Sec17 promotes fusion in either the presence or absence of HOPS — but with faster kinetics when HOPS is absent. Our data further indicate that Sec17 promotes fusion both through its direct penetration of the membrane and by enhancing C-terminal SNARE zippering. In a working model, the interplay among Sec17, Sec18, SMs, and SNARE zippering can explain why SM proteins are indispensable for SNARE-mediated fusion *in vivo*.

**Impact statement:** Sec17 is shown to have divergent effects on pre-fusion SNARE complex activity, depending on the state of SNARE zippering. HOPS, an SM-tether complex, controls the outcome of Sec17-SNARE engagement. The results suggest a coherent working model for SM activity *in vivo*.

## Introduction

Eukaryotic cells partition biosynthetic, catabolic, and information processing activities into specialized organelles. To move materials among organelles while maintaining compartmental identity, specific cargos are picked and packaged into vesicular carriers. These carriers must accurately dock at and fuse with target organelles (Angers and Merz, 2011; Sudhof and Rothman, 2009). Most intracellular fusion events are driven by the concerted folding and oligomerization— zippering — of SNARE proteins embedded, *in trans*, on the two apposed membranes. SNAREs confer inherent compartmental selectivity. Among the many possible permutations of SNARE complexes, only a subset assemble into kinetically stable complexes; among these, only a further subset efficiently initiate fusion (Izawa et al., 2012; McNew et al., 2000).

*Trans*-SNARE zippering, membrane fusion, and *cis*-SNARE disassembly together constitute the SNARE cycle. The folded portion of a SNARE complex is a parallel bundle of α-helical SNARE domains (Qa, Qb, Qc, and R; (Fasshauer et al., 1998; Hanson et al., 1997; Poirier et al., 1998; Sutton et al., 1998). N-to-C zippering of the trans-SNARE complex generates mechanical tension (Hanson et al., 1997; Liu et al., 2006; Min et al., 2013; Zorman et al., 2014), pulling the SNARE C-termini and their associated membranes together to initiate lipid mixing and fusion. After fusion, the SNARE complex membrane anchors lie *in cis*, adjacent and parallel within the membrane. *Cis*-complexes are disassembled to recycle and energize the SNAREs for subsequent fusion reactions (Hanson et al., 1997; Mayer et al., 1996). Disassembly is catalyzed by the Sec18 ATPase, and Sec17, an adapter that docks Sec18 onto the SNARE complex (Hanson et al., 1997; Mayer et al., 1996; Sollner et al., 1993; Zhao et al., 2015). In mammals, Sec18 and Sec17 are named NSF and α-SNAP. Here, we use the yeast nomenclature.

Assembly of the pre-fusion *trans*-SNARE complex is precisely choreographed by tethering and docking factors. Diverse factors operate at specific compartments, but all SNARE-mediated fusion events that have been closely examined require cofactors of the <u>S</u>ec1/<u>M</u>ammalian UNC-18 (SM) family (Baker and Hughson, 2016; Carr and Rizo, 2010; Sudhof and Rothman, 2009). *In vitro*, SM proteins increase the rate of fusion several-fold above the basal rate catalyzed by SNAREs alone. *In vivo*, however, SM deletion annihilates fusion (Carr and Rizo, 2010; Sudhof and Rothman, 2009). It has been unclear whether SM proteins increase the rate or precision of SNARE assembly, make already-formed *trans*-complexes more fusogenic, or both.

Here, we examine SM function in the context of the HOPS tethering complex, which controls endolysosomal docking and fusion (Angers and Merz, 2011). Importantly, the HOPS SM subunit Vps33 is both necessary and sufficient for HOPS interaction with SNARE domains and SNARE complexes (Lobingier et al., 2014; Lobingier and Merz, 2012). A pair of breakthrough crystal structures seems to reveal how the Qa- and R-SNARE interact with Vps33 (Baker et al., 2015). These structures strongly support the hypothesis that Vps33 nucleates and guides assembly of the partially-zipped *trans*-SNARE complex, through a stepwise N-to-C-terminal assembly pathway. The same mechanism may be used by other SM proteins as well.

Perhaps surprisingly, the disassembly adapter Sec17 interacts not only with postfusion SNARE complexes, but with pre-fusion SNARE complexes as well (Barszczewski et al., 2008; Park et al., 2014; Schwartz and Merz, 2009; Wang et al., 2000; Xu et al., 2010; Zick et al., 2015). In an apparent paradox, Sec17 is reported to both inhibit and stimulate fusion. The direct fusogenic activity of Sec17 was first observed in cell-free assays with intact yeast vacuoles (Schwartz and Merz, 2009). To explain these results, we proposed that Sec17 binding to partially-zipped trans-SNARE complexes increases the folding stability of the *trans*-complex’s membrane-proximal C-terminal domain (Schwartz and Merz, 2009). Very recently, single-molecule force spectroscopy experiments provided direct evidence for this zipper-clamp mechanism (Ma et al., 2016). In addition, membrane penetration by a hydrophobic loop near the Sec17 N-terminus is critical for Sec17’s fusogenic activity (Wickner et al; Zick et al., 2015).

We recently discovered that Sec17 and SM proteins can physically associate with one another on quaternary SNARE complexes (Lobingier et al., 2014). However, the functional interplay among Sec17, SMs, and the zippering *trans* complex has not yet been explored, either *in vitro* or *in vivo*. It has been unclear why Sec17 both inhibits and stimulates fusion, and it is unknown how these divergent activities might be regulated. Moreover, it is unknown whether Sec17 can augment the fusion activity of *trans*-SNARE complexes *in vivo* as it does *in vitro*. Here we report parallel studies using chemically-defined proteoliposomes and intact vacuoles *in vitro*, and mutational analyses *in vivo*, to show that SNARE zippering can be precisely manipulated to yield on-pathway trans-complexes that exhibit a range of fusogenic activities. We show that Sec17 functionally interacts with partially-zipped *trans*-complexes during docking — before, as well as after fusion — and demonstrate that the functional consequences of Sec17-SNARE interactions are in turn controlled by HOPS and Vps33. We propose a working model in which the network of physical and functional interactions among Sec17, Sec18, SNAREs, and SM’s explains the absolute *in vivo* requirement for SM proteins in membrane fusion.

## Results

### Qc zippering beyond layer +5 drives fusion *in vivo*

The Qc-SNARE Vam7 is essential for fusion of yeast lysosomal vacuoles. Vam7 is soluble and lacks a transmembrane anchor (Cheever et al., 2001, #431; Sato et al., 1998, #974). Previously, we studied Vam7 C-terminal truncation mutants *in vitro* (Qc-Δ proteins; Fig. 1). A subset of the Qc-Δ mutants formed partially zipped, stalled *trans*-SNARE complexes (Schwartz and Merz, 2009). To test whether Qc-Δ function *in vivo* is like that *in vitro*, we studied cells expressing representative Qc-Δ truncation mutants.

**Figure 1.**
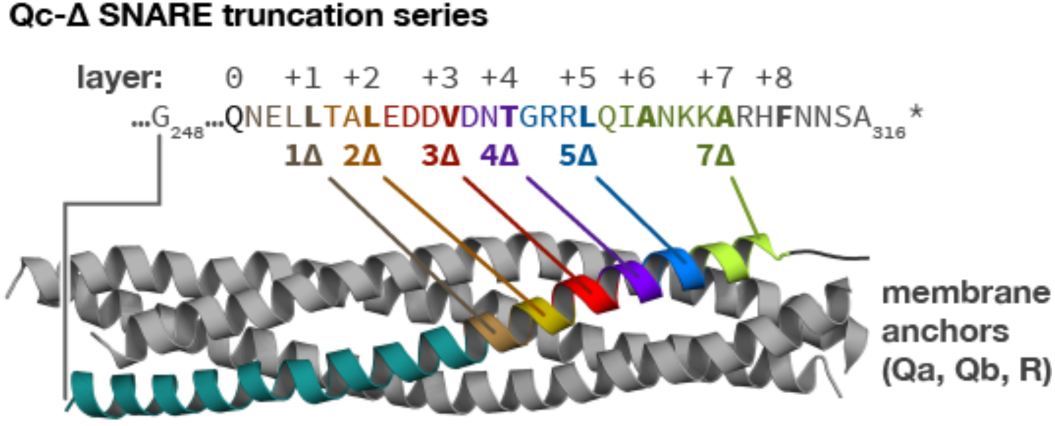
The set of Qc-Δ truncation mutants, mapped onto the structure of a quaternary SNARE bundle. The transmembrane anchors of the Qa-, Qb-, and R-SNAREs are not depicted.

The Qc-Δ mutants were introduced through allelic exchange at the chromosomal *VAM7* locus, or were overproduced from high-copy plasmids. Wild-type yeast cells have 1-5 large vacuoles (class A morphology). Fusion-defective mutants such as *vam7*Δ have numerous fragmented vacuoles (class B; (Raymond et al., 1992; Wada and Anraku, 1992). Consistent with previous *in vitro* experiments (Schwartz and Merz, 2009), Qc-wt or Qc-7Δ knock-ins yielded normal morphology (Fig. 2A), while Qc1Δ, Qc3Δ, Qc-5Δ knock-ins had fragmented vacuoles and were indistinguishable from the *vam7*Δ null mutant. Overproduction of Qc-1Δ, -3Δ, or -5Δ in a *vam7*Δ genetic background also resulted in complete vacuolar fragmentation (Fig. 2B, top row). *In vitro*, Qc3Δ and Qc5Δ nucleate stalled *trans*-SNARE complexes, while Qc1Δ does not enter into stable *trans*-SNARE complexes (Schwartz and Merz, 2009). Consistent with these findings, in wild-type *VAM7* cells (Fig. 2B, bottom row) overproduction of Qc-3Δ or Qc-5Δ, but not Qc1Δ, caused dominant fragmentation with 30-40% penetrance. The partially penetrant phenotype likely reflects cell-to-cell variation in plasmid copy number.

**Figure 2.**
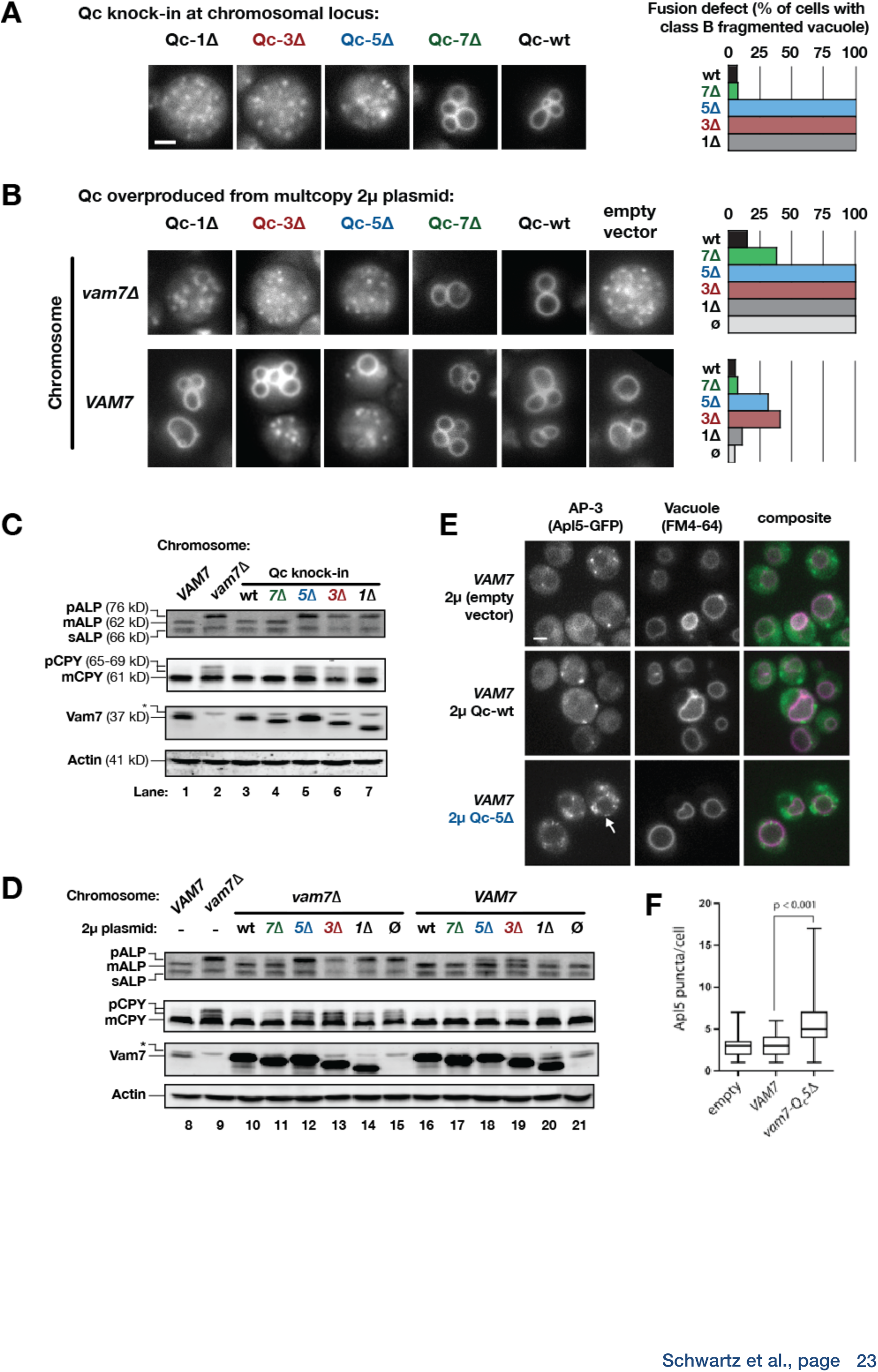
Phenotypic characterization of Qc-Δ mutants ***in vivo*.** A and B. Vacuoles were labeled by pulse-chase with the styryl dye FM4-64 and observed by wide-field epifluorescence microscopy. Defects in vacuole morphology are quantified in the graphs to the right. 106-411 cells of each genotype were scored in at least two independent experiments. **C** and **D**. Cargo trafficking defects of Qc-Δ mutants. Cell lysates were separated by SDS-PAGE and analyzed by immunoblot using antibodies against Vam7 (Qc), ALP, or CPY. Anti-actin is used for loading controls. The aberrant migration pattern of Qc-5Δ is also seen with recombinant Qc-5Δ purified from *E. coli* cells. A non-specific band present in all strains including *vam7*Δ mutants is indicated by (*). **E and F**. Overproduction of Qc5Δ causes dominant partial accumulation of AP-3 vesicles. In some cells, the vacuole is not fragmented and AP-3 vesicles accumulate at the vacuole limiting membrane, as shown in E. In F, AP-3 vesicles are quantified (Mann-Witney *U* test; n = 100 cell profiles per strain in two independent experiments). Scale bars (A,B,E) indicate 2μm.

In addition to homotypic fusion, Vam7 is needed for heterotypic fusion of vesicular carriers with the vacuole. We examined two pathways. Alkaline phosphatase (ALP) traffics directly from Golgi to vacuole, while carboxypeptidase Y (CPY) traffics from the Golgi to the vacuole *via* late endosomes. ALP and CPY traffic as inactive proenzymes that are processed and activated at the vacuole. Fusion defects cause slow-migrating pro-forms, pALP and pCPY, to accumulate. Cells expressing Qc-1Δ, -3Δ, or -5Δ as knock-ins at the chromosomal *VAM7* locus had ALP and CPY maturation defects as severe as the *vam7*Δ null mutant (Fig. 2C, compare lane 2 to lanes 5-7). Comparable defects were observed when Qc-1Δ, -3Δ, or -5Δ were overproduced from high-copy plasmids in the *vam7*Δ null background (Fig. 2D, lanes 12-14).

In the *VAM7* wild-type background, Qc-3Δ or Qc-5Δ overproduction caused dominant, partial defects in ALP maturation (Fig. 2D, lanes 18 and 19). ALP is carried from the trans-Golgi to the vacuole in vesicles bearing the AP-3 coat complex (Cowles et al., 1997). When docking and fusion at the vacuole are impaired, AP-3 vesicles accumulate (Angers and Merz, 2009; Rehling et al., 1999). In wild-type cells overproducing Qc-5Δ, the median number of AP-3 vesicles nearly doubled (Fig. 2E,F). Qualitatively similar accumulations of AP-3 vesicles were observed in Qc-3Δ overproducers. Moreover, in Qc-5Δ-overproducer cells, AP-3 puncta were observed in clumps at the vacuole limiting membrane (Fig. 2E, arrow), suggestive of a defect in fusion rather than docking, and further implying that dissociation of the AP-3 coat does not normally occur until after the completion of SNARE zippering (Angers and Merz, 2009). These results for homotypic and heterotypic fusion *in vivo* are broadly consistent with our previous analyses of Qc-Δ mutants *in vitro* (Schwartz and Merz, 2009).

### Sec17 interacts with partially-zipped SNAREs to control fusion

To see if we could detect additional functional transitions during SNARE zippering, we characterized two additional truncation mutants, Qc-2Δ and Qc-4Δ using the cell-free assay of vacuole homotypic fusion. This assay employs enzymatic complementation to quantify luminal content mixing between native lysosomal vacuoles (Fig. S1A; (Merz and Wickner, 2004a). In “ATP bypass” gain-of-function assays (Fig. S1B, reaction ii), neither Qc-2Δ nor Qc4Δ drove fusion, even at high concentrations (Fig. 3A). To determine whether these mutants entered *trans*-complexes, like Qc3Δ, or failed to enter trans-complexes, like Qc-1Δ, we assayed competitive inhibition of fusion driven by native Qc-wt (Fig S1B, reaction i). In these experiments Qc-2Δ, like Qc1Δ, was weakly inhibitory, while Qc-4Δ, like Qc-3Δ, was a potent competitive inhibitor (Fig. 3B). We infer that Qc-4Δ, like Qc-3Δ (Schwartz and Merz, 2009; Xu et al., 2010), enters into partially-zipped, fusion-defective *trans*-SNARE complexes. We further propose that this state corresponds to the metastable, partially-zipped SNARE conformations observed in force spectroscopy studies (Liu et al., 2006; Min et al., 2013; Zorman et al., 2014). Indeed, in the “half-zipped” state, the neuronal t/Q-SNARE complex is structured to layer +4 (Ma et al., 2016), a finding in precise agreement with our experiments.

**Figure 3.**
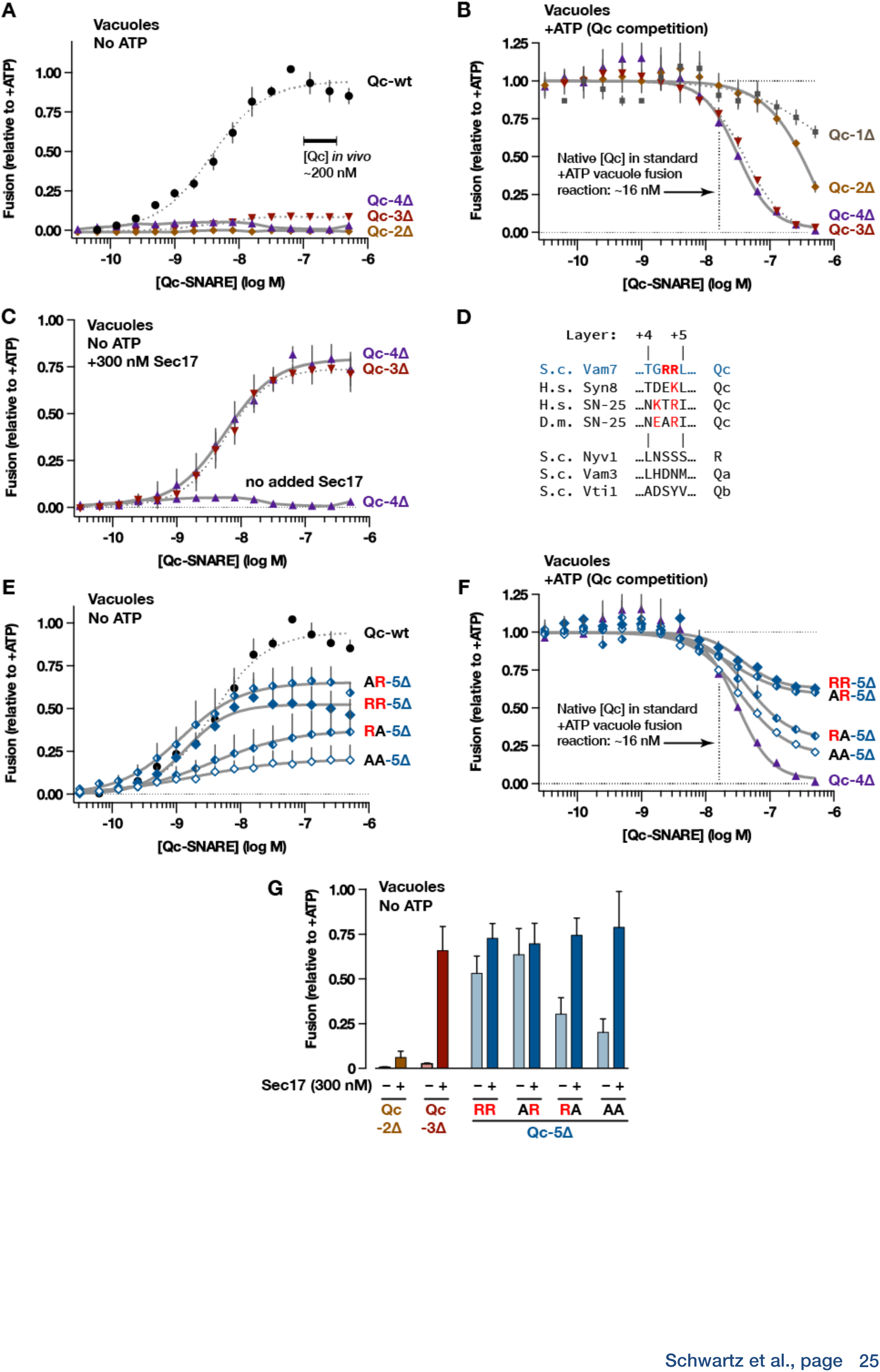
Interplay of Sec17 and SNARE zippering in cell-free assays of homotypic vacuole fusion. The content-mixing assay system and the reaction schemes are diagrammed in Fig. S1. **A**. Qc-2Δ, -3Δ, and -4Δ are nonfusogenic in gain-of function assays. **B**. Qc-3Δ and -4Δ are efficient competitive inhibitors of Qc-wt. **C**. Added Sec17 restores fusion activity to Qc-3Δ and -4Δ in the absence of ATP. **D**. Sequence alignment of SNAREs in the layer +4 to +5 region. S.c., *Saccharomyces cerevisiae;* H.s., *Homo* sapiens; D.m., *Drosophila* melanogaster. A conserved arginyl (R) residue is indicated in red. **E**. Gain-of-function assays with Qc-5Δ variants. **F**. Competitive inhibition of fusion by Qc-5Δ variants. **G**. Sec17 rescue assays for Qc-2Δ, 3Δ, and 5Δ variants The Qc-Δ SNAREs were added to 100 nM. Lines in the dose-response and dose-inhibition experiments show nonlinear fits of Hill equation. Dashed lines denote data re-plotted from Schwartz and Merz (2009) to facilitate comparison. For all panels, points and bars indicate mean (+ or ± s.e.m.) of ≥3 independent experiments.

Sec17 addition rescues the ability of Qc-3Δ to drive fusion in the apparent absence of Sec18 or ATP (Schwartz and Merz, 2009; Wickner et al.). Sec17 rescued Qc-4Δ reactions with a dose-response indistinguishable from that of Qc-3Δ (Fig. 3C). In contrast, Sec17 failed to rescue fusion in reactions containing Qc-2Δ (see Fig. 3G). Taken together these results and our previous studies (Schwartz and Merz, 2009) indicate that the propensity to form kinetically stable *trans* complexes increases in a switch-like manner as the Qc-SNARE assembles from layer +2 to layer +3. The propensity of the partially-zipped *trans*-complex to initiate fusion then sharply increases abruptly with a single α-helical turn, from Qc layer +4 to +5 (Fig. 3C,E), and further increases to layer +7 (Schwartz and Merz, 2009). With a stalled *trans*-complex zipped to Qc layer +3 or +4, the membranes should be separated by ∼8-10 nm.

The sharply increased fusion activity of Qc5Δ versus Qc4Δ prompted closer examination of the layer +5 region. Vam7 contains two arginyl (R) residues between layers +4 and +5 (Fig. 3D). The second residue is conserved in many Qc-SNAREs (Fasshauer et al., 1998; Sutton et al., 1998). In gain-of-function experiments (Fig. 3E), mutation of the non-conserved first Arg to Ala (AR) slightly increased fusion versus the “wild-type”(RR) Qc-5Δ. Mutation of the second, conserved Arg (RA) decreased fusion. Mutation of both residues (AA) further impaired fusion. In competition assays, the abilities of the variants to dominantly impair fusion corresponded to their activities in gain-of function assays (Fig. 3F; compare to 3E). The apparent K_M_ (EC_50_) of the mutant Qc5Δ’s (AR, RA, AA) were indistinguishable from “wild-type” Qc-5Δ (RR), suggesting that the ability of the Qc5Δ mutants to enter into pre-fusion complexes was unaltered, even as their capacity to drive fusion diverged. Moreover, added Sec17 allowed all Qc-5Δ variants to drive fusion with comparable efficiency (Fig. 3G). These findings, and results from other systems (Fasshauer et al., 1998; Mohrmann et al., 2010; Sakaba et al., 2005) indicate that Qc layer +5 has an absolutely pivotal role in fusion, probably driving SNARE assembly beyond the “half-zipped” metastable state (Liu et al., 2006; Ma et al., 2016; Min et al., 2013; Zorman et al., 2014). Again, our results with intact vacuoles agree precisely with force spectroscopy studies, which show that Sec17 (α-SNAP) stabilizes the folded state of complexes that are zipped past layer +4 (Ma et al., 2016).

### Sec17 rescues partially-zipped complexes in the complete absence of Sec18

To establish the minimal requirements for Sec17 rescue of stalled *trans-*complexes, we used chemically-defined reconstituted proteoliposomes (RPLs; Fig. S2; Zick et al., 2015; Wickner et al; Zick et al., 2014; Zucchi and Zick, 2011). To bypass requirements for ATP and Sec18 (Mayer et al., 1996; Schwartz and Merz, 2009; Thorngren et al., 2004), the SNAREs were distributed asymmetrically, as in the heterotypic fusion configuration. One set of RPLs displayed the Qa- and Qb-SNAREs Vam3 and Vti1; the other, the R-SNARE Nyv1 (Fukuda et al., 2000). All RPLs displayed GTP-bound Ypt7 Rab. Lipid mixing and luminal aqueous content mixing were simultaneously monitored in each reaction using Förster resonance energy transfer (FRET) probes (see Fig. S2). Both read-outs yielded similar results. We therefore focus on content mixing, the reaction endpoint.

When HOPS and full-length Qc-wt were added to RPLs, fusion was rapid and efficient, with or without Sec17 (Fig. 4A-C). In contrast to Qc-wt, the truncation mutants Qc-3Δ, -4Δ, and -5Δ were non-fusogenic until Sec17 was supplied (Fig. 4B,C). In the absence of Sec17, Sec18 was unable to rescue Qc truncation mutants. However, Sec18 dramatically enhanced Sec17-mediated rescue and allowed rescue at much lower Sec17 concentrations (Fig. 4D-I). In a companion study (Wickner et al.) we demonstrate that Sec18 can augment Sec17-mediated Qc-3Δ rescue even in the absence of ATP hydrolysis. We conclude that Sec17 alone, or Sec17 and Sec18 together, can stimulate fusion independently of Sec18-catalyzed SNARE complex disassembly.

**Figure 4.**
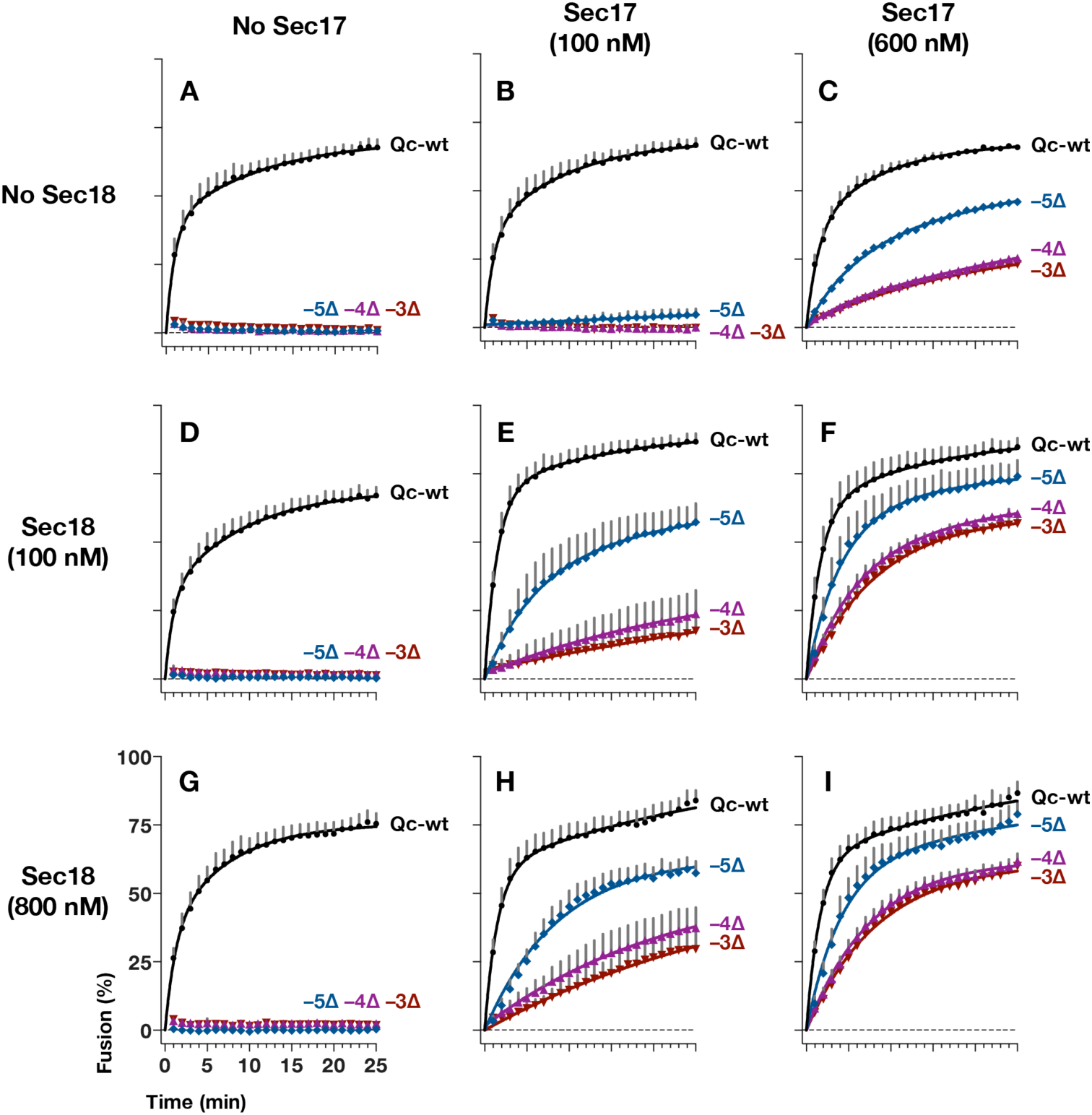
Sec17 allows partially-zipped SNARE complexes to drive fusion with or without Sec18. The chemically-defined RPL fusion system is diagrammed in Fig. S2. RPLs bearing Ypt7-GTP (Rab), and either Qa- and Q-b, or R-SNAREs, were incubated with HOPS (100 nM) and the indicated Qc-SNAREs (250 nM), and in the absence or presence of Sec17 and Sec18 as indicated. Each data point shows the mean content mixing signal + s.e.m. for 3 independent experiments. The lines show nonlinear best-fits of a second-order kinetic model.

These experiments allow us to draw several important conclusions. First, Sec17 rescue of partially-zipped SNARE complexes requires neither Sec18 nor ATP hydrolysis, consistent with the vacuole experiments as well as our more recent experiments using RPLs and a Qb-SNARE lacking its transmembrane segment (Zick et al., 2015). Second, Sec17-mediated rescue does not entail disassembly of post-fusion *cis*-complexes. Again, our work with RPLs and a different truncated SNARE, Qb-ΔTMD, further buttresses these conclusions (Zick et al., 2015; Wickner et al.). Third, native Vam7 (Qc-wt) might be available in residual quantities on unprimed vacuoles, but it is not present in the Qc-Δ RPL reactions. Thus, Sec17 supports Qc-Δ–mediated fusion in the total absence of full-length Qc-wt. Fourth, no additional vacuolar proteins are needed for Sec17 stimulation of fusion. These findings and other results (Ma et al., 2016; Park et al., 2014; Schwartz and Merz, 2009; Xu et al., 2010; Zick et al., 2015), definitively establish that Sec17 and Sec18 interact not only with *post*-fusion *cis*-complexes to mediate SNARE disassembly, but both physically and functionally with *pre*-fusion *trans*-SNARE complexes in various states of zippering — at least, *in vitro*.

### Sec17 augments Qc-Δ SNARE function *in vivo*

Sec17 addition allows Qc-3Δ, -4Δ, or -5Δ to fuse vacuoles and RPLs (Figs. 3 and 4). However, all Qc-Δ’s but Qc-7Δ had severe fusion defects *in vivo*, where Sec17 and Sec18 are active (Fig. 3). We therefore asked whether overproduction of Sec17, Sec18, or both together might suppress the functional defects of QcΔ SNAREs *in vivo*. To verify that overproduced Sec17 and Sec18 are functional *in vivo*, we assayed SNARE complex abundance by co-immunoprecipitation (Fig. 5A). At steady state only ∼3% of SNARE complexes are *in trans*, so this approach assays the ∼97% of complexes *in cis*. Sec18 overproduction, with or without Sec17 overproduction, decreased *cis*-complex abundance and decreased Sec17-SNARE association (Fig. 5A, compare lanes 2,3 to 4,5). In contrast, Sec17 overproduction alone did not substantially alter either *cis*-complex abundance or Sec17-SNARE association (Fig. 5, compare lanes 2 and 3). Similar results were previously obtained with *Drosophila* (Babcock et al., 2004; Golby et al., 2001).

**Figure 5.**
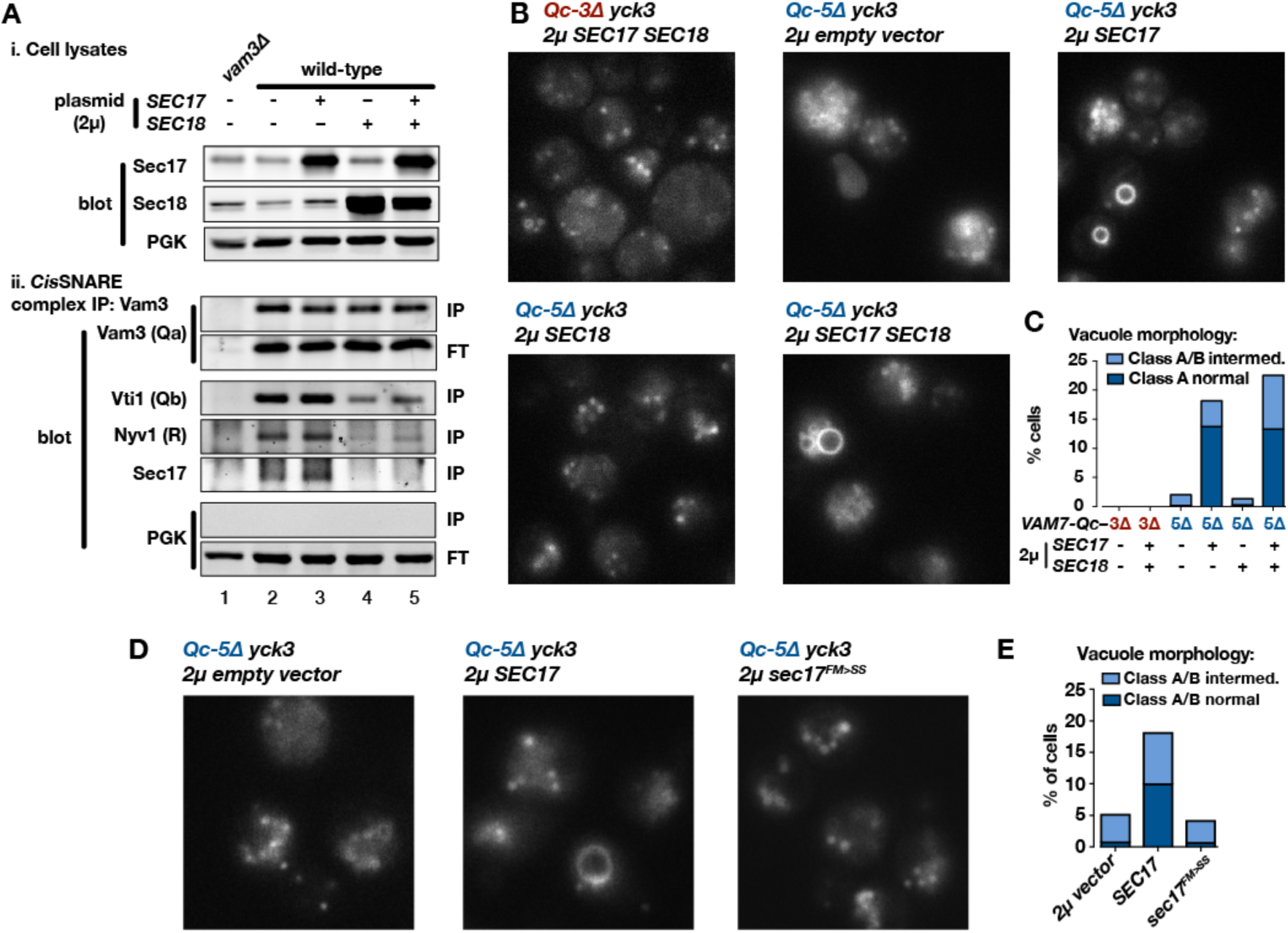
Sec17 overproduction partially rescues ***in vivo*** activity of Qc-5Δ and depends on the Sec17 hydrophobic loop. **A**. Analysis of *cis*-SNARE complex abundance in lysates of cells overproducing Sec17 and Sec18. Anti-Vam3 (Qa-SNARE) immunoprecipitates from lysates of the indicated strains were separated by SDS-PAGE and analyzed by immunoblot as indicated. IP, immunoprecipitate; FT, flow-through. PGK, phosphoglycerate kinase (control). **B**. Vacuoles were labeled with FM4-64 dye and observed by epifluorescence microscopy. **C**. Quantification of phenotypes in B. Bars show mean scores from three independent experiments (n=88-354 cells per genotype per experiment). **D and E**. Sec17 with a less-hydrophobic N-terminal loop fails to rescue Qc-5Δ function in vivo.

Overproduction of Sec17, Sec18, or both together did not alter vacuole morphology in otherwise wild-type cells, and failed to rescue vacuole morphology in cells expressing Qc-3Δ or Qc-5Δ (Fig. S3). There is, however, an important difference between the conditions *in vivo* and *in vitro. In vivo*, the vacuole-associated kinase Yck3 negatively regulates fusion by phosphorylating HOPS and the Rab GEF Mon1. These phosphorylation events impair both Ypt7 Rab activation and HOPS-membrane association (Brett et al., 2008; LaGrassa and Ungermann, 2005; Lawrence et al., 2014). The Sec17 rescue reactions with purified yeast vacuoles lacked ATP, resulting in Yck3 loss-of-function (Brett et al., 2008), and Yck3 was not present in the RPL reactions. Thus, we tested Qc-Δ function in *yck3*Δ mutant cells. In *yck3*Δ cells expressing either Qc-3Δ or Qc-5Δ, vacuoles were uniformly fragmented, indicating that elevated HOPS activity does not by itself restore Qc-Δ function. However, overproduction of either Sec17, or Sec17 and Sec18 together, rescued Qc-5Δ vacuole morphology with ∼20% penetrance (Fig. 5B,C). The partial rescue of Qc-5Δ rescue makes sense. In the cell-free assay of homotypic vacuole fusion, Sec17 restores Qc-Δ activity only over a specific range of Sec17 concentrations (Fig. S4A). Because the copy number of yeast 2μ plasmids varies from cell to cell, partial rescue probably reflects variation in Sec17 expression. Qc-3Δ mutants exhibited severe defects *in vivo* under all conditions, consistent with the generally lower fusion activity of Qc-3Δ *in vitro*. Sec17 overproduction alone had no influence on *cis-* SNARE complex abundance, and Sec18 overproduction alone did not rescue Qc-5Δ (Fig. 5B,C). This means that neither the rate of *cis*-SNARE complex disassembly, nor the steady-state availability of unpaired SNAREs, can explain why Sec17 augments Qc-5Δ function *in vivo*. The most likely remaining explanation is that Sec17 rescues partially-zipped *trans*-complexes *in vivo* just as it does *in vitro*.

Sec17 and mammalian α-SNAP contain a flexible N-terminal loop bearing four hydrophobic residues that penetrate the phospholipid bilayer (Winter et al., 2009). Mutation of two residues (Sec17^FM>SS^; see Fig. S4B) nearly eliminated Sec17 stimulation of fusion in both RPLs (Zick et al., 2015; Wickner et al.) and cell-free homotypic fusion assays (Fig. S4A). Similarly, Sec17^FM>SS^ failed to rescue vacuole morphology in *yck3*Δ Qc*-5*Δ cells (Fig. 5D,E). We conclude that membrane penetration by Sec17 is central to its ability to stimulate fusion — not only with synthetic RPLs but with intact vacuoles, both *in vitro* and *in vivo*.

### Sec17 stimulates fusion after docking is complete

Where in the forward docking and fusion pathway does Sec17 act to stimulate fusion? To begin to address this question we did staging experiments with intact vacuoles. Vacuoles were pre-incubated for 10 min to allow initial association (tethering). Then either buffer (control condition) or Qc3Δ was added. The vacuoles were further incubated for 15 min to allow the Qc-3Δ reactions to assemble *trans*-SNARE complexes. Following this incubation, fusion was triggered by adding either full-length Qc-wt to the control reaction, or Sec17 to the Qc3Δ reaction. In the control reaction (Fig. 6A), stable *trans-*SNARE complexes do not assemble until Qc-wt is added (Boeddinghaus et al., 2002; Schwartz and Merz, 2009; Thorngren et al., 2004), whereas in the Qc-3Δ reaction, *trans*-SNARE complexes form during the 15 min prior to Sec17 addition (Schwartz and Merz, 2009). As in previous experiments (Boeddinghaus et al., 2002; Merz and Wickner, 2004b; Thorngren et al., 2004), the control reaction was potently inhibited by antibody against the Qa-SNARE Vam3, even when the antibody was added only 1 min before Qc-wt. The control reaction was also inhibited by anti-Ypt7 (yeast Rab7) and anti-Vps33 (the SM subunit of HOPS), up to the –1 min time point. Thus, as expected, docking factors are susceptible to inhibition at least until the onset of *trans*-SNARE complex assembly.

**Figure 6.**
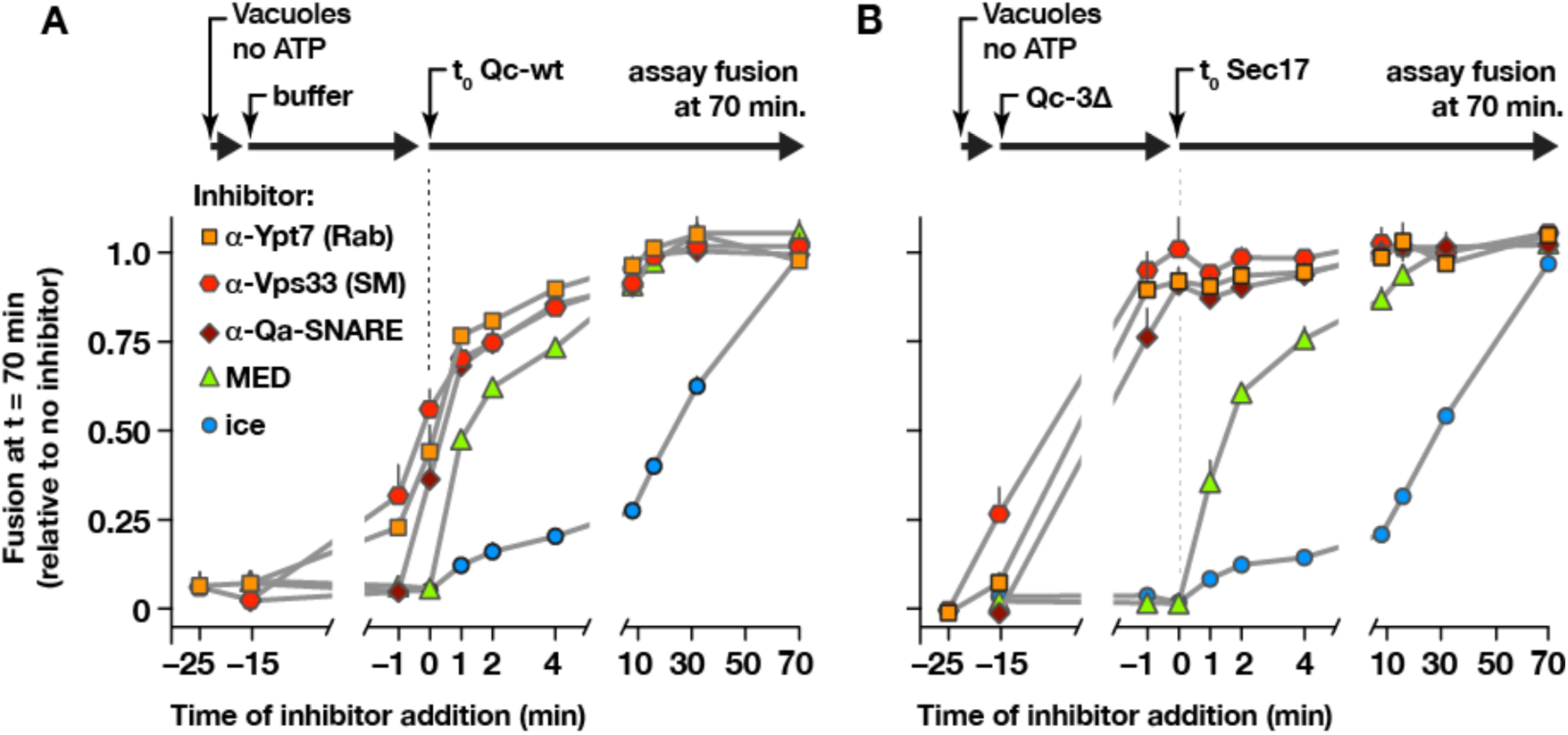
Staging of Sec17 rescue in cell-free assays of vacuole fusion. A. Master fusion reactions were pre-incubated for 25 min at 30° C, and fusion was initiated by adding Qc-wt (20 nM). **B**. Master fusion reactions were pre-incubated for 25 min at 30° C in the presence of 300 nM Qc-3Δ, and fusion was initiated by adding Sec17 (300 nM). For both A and B, at the indicated times single-reaction aliquots were removed from the master reactions, and combined with inhibitors as specified (or moved to ice). For all reactions, fusion was assayed at the 70 min. time point. Points indicate mean + s.e.m. for 2-6 independent experiments.

In the Qc-3Δ / Sec17 rescue experiment (Fig. 6B), addition of anti-Ypt7 (Rab), anti-Vps33 (SM), or anti-Vam3 (Qa-SNARE) prior to Qc3Δ addition eliminated Sec17-triggered fusion. However, during the pre-Sec17 incubation, Qc-3Δ reactions gained resistance to all docking inhibitors. By –1 min prior to Sec17 addition, and at subsequent time points, the reaction was insensitive to antibodies against the Rab, the HOPS SM subunit Vps33, or the Qa-SNARE Vam3. Qc-3Δ addition therefore drives the vacuoles into an operationally docked state. In this state kinetically stable *trans*-SNARE complexes have assembled (Schwartz and Merz, 2009; Xu et al., 2010), and Sec17 efficiently triggers fusion even when inhibitors of the Rab, the SM, or the Qa-SNARE are present. A post-docking inhibitor of fusion, the membrane-interacting MARCKS effector domain (MED), inhibited both reaction schemes (Fig. 6A,B) up to the moment of Sec17 or Qc-wt addition. These findings imply that Sec17 need not be present during tethering or docking, but rather that it acts to trigger fusion once docking is complete. Moreover, the results imply, — but do not prove — that Sec17 might be able to trigger fusion by partially-zipped SNARE complexes in the absence of HOPS and its SM subunit. To test these ideas under conditions permitting greater experimental control, we returned to the chemically defined RPL system.

### HOPS selects the outcome of Sec17–SNARE interactions

HOPS promotes efficient tethering and docking (Stroupe et al., 2009; Zick et al., 2014). To test whether HOPS influences the function of Sec17 during docking and fusion, parallel RPL reactions were initiated with Qc-wt in the absence or presence of Sec17, and in the absence or presence of HOPS. In the absence of HOPS, vesicle tethering and *trans*-SNARE assembly can be driven by 2% polyethylene glycol (PEG; (Hickey and Wickner, 2010; Zick et al., 2014). In control reactions full-length Qc-wt drove efficient fusion in the presence of HOPS, and this fusion was unaffected by Sec 17 (Fig. 7A). In marked contrast, in no-HOPS reactions (Fig. 7B) Sec17 strongly impaired fusion driven by Qc-wt. Inhibition of fusion *in vitro* by Sec17, or by Sec17 and Sec18 together, has been reported in multiple systems (Ma et al., 2013; Park et al., 2014; Wang et al., 2000; Xu et al., 2010). Moreover, overproduced Sec17 is toxic *in vivo* when SM (Vps33 or Sly1) function is partially impaired (Lobingier et al., 2014). These findings suggest that HOPS allows fusion to occur in the presence of otherwise inhibitory Sec17.

**Figure 7.**
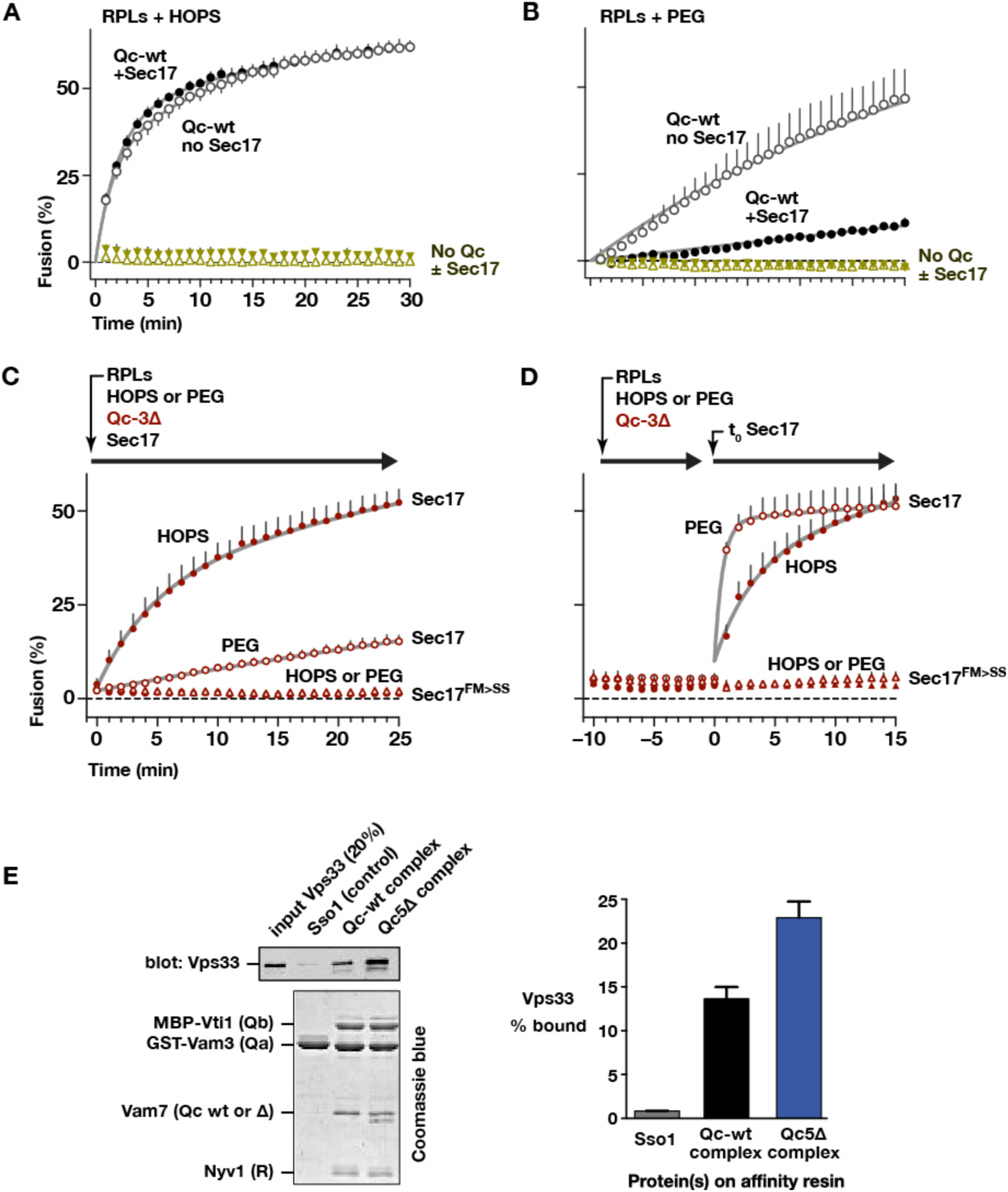
Staging experiments show that HOPS-SM gates Sec17 function. A and B, reactions were initiated as in Fig. 2, except that HOPS was either present (100 nM) or replaced by 2% PEG. Sec17 was present at 0 or 600 nM, and Vam7 was present at either 80 or 400 nM in panel A or B, respectively. **C** and **D**. Staging of Sec17 effects. **C**. Simultaneous addition of Qc-3Δ, Sec17 or Sec17^FM>SS^ (600 nM), and HOPS (50 nM) or PEG (2%). **D**. As in E, except that Sec17 or Sec17^FM>SS^ was added following a 10 min pre-incubation at 27° C. For A-D points denote mean + s.e.m. of 2 (A,B) or 6 (B,C) independent experiments. Lines show nonlinear best-fits of a second-order kinetic model.

How can Sec17 inhibition of fusion in the absence of HOPS be reconciled with Sec17 triggering of fusion in the presence of HOPS? The staging experiments with vacuoles (Fig. 6) implied that HOPS and its SM subunit might be dispensable once *trans*-SNARE complexes have formed. This hypothesis predicts that Sec17 has different functions before and after *trans*-SNARE complex zippering. We therefore tested whether Sec17 can trigger fusion by stalled Qc-Δ *trans*-complexes established in the complete absence of Sec17 and HOPS. In reactions where the Qc-Δ and Sec17 were added together, (Fig. 7C), Sec17 stimulated Qc-3Δ-driven fusion in the presence of HOPS, and far less efficiently without HOPS. As expected, Sec17^FM>SS^ was completely unable to promote fusion, verifying that Sec17 membrane penetration is required for Sec17-triggered fusion. Next, staged reactions were initiated with Qc-3Δ in the presence of PEG or HOPS. These reactions were incubated for 10 min to allow *trans*-SNARE complexes to pre-assemble (Fig. 7D). Sec17 was then added to trigger fusion. Remarkably, Sec17 triggered fusion by pre-formed Qc-3Δ complexes in the PEG reactions just as efficiently as in the HOPS reactions. Thus, HOPS not only accelerates tethering and productive *trans*-SNARE zippering, but allows these processes to occur in the presence of otherwise-inhibitory Sec17. Once the partially-zipped SNARE complex has assembled, HOPS (and its SM subunit Vps33) are dispensable. Indeed, the SM may need to dissociate from the *trans*-complex for terminal zippering and membrane fusion to occur, a hypothesis strongly supported by recent Vps33-SNARE crystal structures (Baker et al., 2015). As predicted by this model, Sec17 triggers fusion with ∼5-fold faster kinetics in the absence versus the presence of Vps33 and HOPS (Fig. 5F). An additional prediction is that purified Vps33 might bind more efficiently to quaternary Qc-Δ SNARE complexes with splayed C-termini. In solution-phase binding assays this was indeed the case (Fig. 7E).

## Discussion

Deletion of SM proteins *in vivo* usually causes an absolute fusion defect. Hence, SMs are considered part of the core SNARE fusion machinery (Carr and Rizo, 2010; Sudhof and Rothman, 2009). However, the mechanisms through which SMs promote fusion have been unclear. Some SMs stimulate SNARE-mediated fusion *in vitro* by several-fold in the absence of Sec17 and Sec18, but this activity is probably not sufficient to account for the absolute SM requirements consistently observed *in vivo*. In the presence of Sec17 and Sec18, however, SM activity is indispensible for fusion, both *in vitro* (Ma et al., 2013; Mima et al., 2008) and *in vivo* (Lobingier et al., 2014). The present experiments demonstrate that Sec17 can interact with zippering SNAREs independently of Sec18 or Sec18 catalytic activity, and reveal that the outcome of the Sec17-SNARE interaction is gated by HOPS and its SM subunit, Vps33 (Fig. 8A).

**Figure 8.**
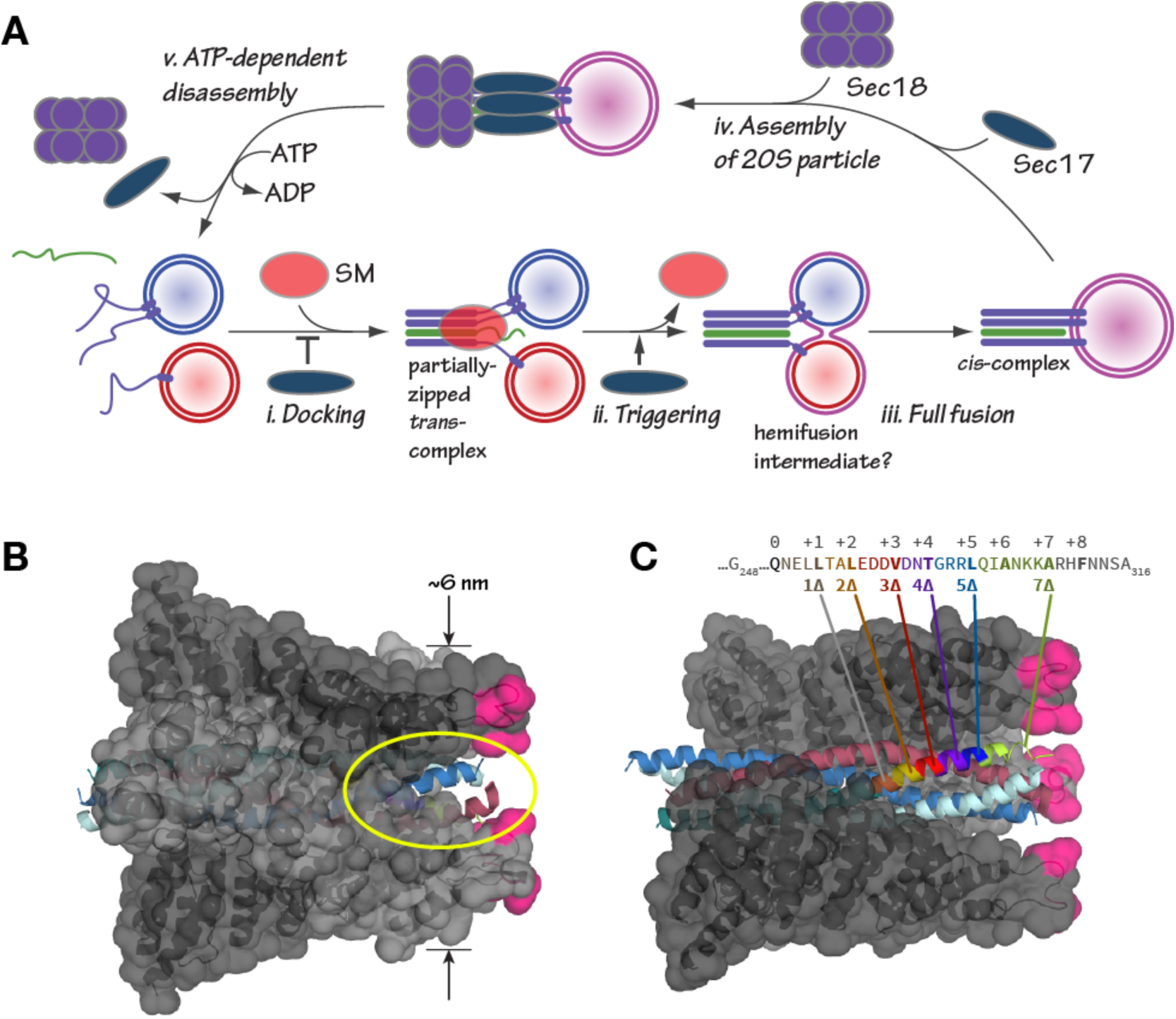
**A**, Working model: interplay of Sec17 and SM-tethering complex on forward fusion pathway. See Discussion for details. **B** and **C**, Sec17/a-SNAP on quaternary SNARE complex (renderings based on PDB 3J96; (Zhao et al., 2015). **In B, note the tapered overall shape of the complex, which forms a wedge-like structure that could fit between two tightly-docked membranes. The hydrophobic loop is magenta. The yellow ellipse shows one of four portals between adjacent Sec17 molecules, through which unstructured SNARE juxta-membrane regions could link the partially-zipped *trans*-SNARE complex to the SNARE transmembrane helices. In C, one of four Sec17 molecules is removed to reveal the Qc packing layers. The portal openings begin at layer +6. Sec17 partially contacts SNAREs to layer +7 or +8.**

Accumulating evidence shows that Sec17 operates not only downstream of fusion to recruit Sec18 but upstream, during docking and SNARE assembly. First, Sec17 physically associates with *trans*-SNARE complexes (Zick et al., 2015; Park et al., 2014; Xu et al., 2010). Second, Sec17 can inhibit SNARE-mediated fusion by acting either prior to SNARE zippering or at an early stage of zippering (Fig. 5; (Park et al., 2014; Wang et al., 2000). Third, Sec17 can trigger fusion by interacting with partially-zipped SNARE complexes (this study; (Zick et al., 2015; Schwartz and Merz, 2009). Fourth, Sec17 and Sec18 together can disassemble stalled *trans*-SNARE complexes. (Rohde et al., 2003; Xu et al., 2010). Fifth, Sec17 and SM proteins can simultaneously and cooperatively bind quaternary SNARE bundles (Lobingier et al., 2014).

Recent structures of mammalian SNARE-Sec17-Sec18 (20S) particles reveal a Sec17 geometry that is precisely optimized for interaction with both *cis-* and partially-zipped *trans*-complexes (Fig. 8B). Up to 4 Sec17 molecules bind the core SNARE complex (Zhao et al., 2015), and up to three Sec17 molecules can bind the core SNARE complex when an SM protein is also bound (Lobingier et al., 2014). Consistent with these findings, Sec17 rescue of stalled Qc-Δ complexes is highly cooperative (Schwartz and Merz, 2009). Membrane bilayers bridged by a “half-zipped” *trans*-complex are 8-10 nm apart (Liu et al., 2006; Min et al., 2013; Zorman et al., 2014). The membrane-proximal end of the wedge-shaped Sec17-SNARE complex is ∼6 nm in diameter (Zhao et al., 2015).

Membrane penetration by the N-terminal hydrophobic loop of Sec17 increases Sec17 affinity for membrane-bound SNARE complexes and is required for Sec17 triggering of fusion, as shown here and in other recent work (Wickner et al; Zick et al., 2015). The loop is disordered in Sec17 crystals, and it is likely to be very flexible (Rice and Brunger, 1999). Between each pair of membrane-penetration loops is an open portal (Zhao et al., 2015). The portals would allow the SNARE residues linking the partially-zipped helical bundle to its transmembrane anchors to pass cleanly between adjacent Sec17 subunits.

Experiments with vacuoles (Schwartz and Merz, 2009) and RPLs (AJM, unpublished results) show that Sec17 promotes the onset of lipid mixing, not the resolution of pre-formed hemifusion intermediates. How then does Sec17 promote fusion? First, Sec17 could stabilize the partially-zipped SNARE complex. Consistent with this idea, Sec17 contacts the Qc precisely up to layer +5. Second, Sec17 bound to the trans complex may operate as a wedge that increases local membrane bending energy at the fusion site. Third, the hydrophobic residues in Sec17 may locally perturb the outer bilayer leaflet, as proposed for synaptotagmin in Ca^2+^-triggering of synchronous neurotransmitter release. Indeed, Sec17 and synaptotagmin compete for binding to SNARE complexes (Sollner et al., 1993), suggesting that Sec17 and synaptotagmin are alternative factors that execute analogous triggering functions. Fourth, we have previously shown that three Sec17 molecules and one SM can bind to a single SNARE complex (Lobingier et al., 2014). A fourth Sec17 molecule might displace the SM, allowing for the completion of SNARE zippering. The hypothesis that multiple Sec17 molecules engage a *trans*-SNARE complex to promote fusion is supported both by the strong cooperativity of Sec17 activity (Schwartz and Merz, 2009), and by the fact that rigor-locked, ATPץS-bound Sec18 augments the fusogenic activity of Sec17 (Wickner et al.).

It is increasingly clear that SM proteins have the properties of true enzymes. In our working model (Fig. S6) SM proteins promote assembly of the *trans*-SNARE complex to its metastable half-zipped state, while preventing non-productive off-pathway reactions, particularly early inhibition of fusion by Sec17. In addition, the SNARE-bound SM prevents premature SNARE complex disassembly by Sec17 and Sec18 (Lobingier et al., 2014; Ma et al., 2013; Xu et al., 2010). Finally, the SM probably inspects the composition and shape of the incipient *trans*-SNARE complex; if the SM fails to bind, or if the SM dissociates, the inhibitory activities of Sec17 and Sec18 are unrestrained: fusion is blocked and the complex is disassembled. Tests *in vivo* support key predictions of this model. First, Sec17 overproduction rescues fusion driven by Qc-5Δ *in vivo*. Second, this rescue occurs only when HOPS/SM activity is augmented. These results suggest that the basal level of *in vivo* HOPS activity is insufficient to overcome Sec17 inhibition of early docking when Sec17 is overproduced. Third, wild-type cells show no overt defects when Sec17 is overproduced, but Sec17 causes trafficking defects and toxicity in cells with partial SM deficiency (Lobingier et al., 2014). The overall picture that emerges is one where SM proteins functionally interact with SNAREs and Sec17 during docking, and SMs dictate the functional outcome of the Sec17-SNARE interaction.

## Materials & Methods

### Plasmids and yeast strains

Yeast culture and genetic manipulations were done using standard methods (Amberg et al., 2005). Strains and plasmids are listed in Supplementary Table S1. *E. coli* expression vectors pMS108-12 were constructed as described (Schwartz and Merz, 2009), with mutations encoded in the primers used to amplify the *VAM7* sequence. The *VAM7* knock-in plasmid pMS125 was generated by ligating two homology arms encompassing the entire *VAM7* coding sequence and surrounding regulatory sequences on either side of the *NAT1* cassette in pAG25 (Goldstein and McCusker, 1999). pMS126-9 were generated by oligo-directed mutagenesis of pMS125 to insert early stop codons. *VAM7* chromosomal knock-ins were constructed by homologous recombination in *vam7*Δ*::KAN* yeast directed by linearized pMS125-9 followed by selection on nourseothricin. pMS120 was constructed by cloning a 2.1 kb fragment of yeast genomic DNA encompassing the entire *VAM7* coding sequence and surrounding regulatory regions (from -1000 to +1210 relative to the translational start site) into pRS426. pMS121-4 were generated by oligonucleotide-directed mutagenesis of pMS120 to introduce early stop codons. *Chromosomal YCK3* loci were ablated via homologous recombination with a PCR product derived from *yck3Δ* null cells (*yck3*Δ*::KAN*). pDN368 was generated by sequence overlap extension PCR of a *SEC17* template to introduce F21S/M22S point mutations, followed by gap repair cloning of the resulting PCR product into SacI-digested pDN526.

### Proteins and lipids

Sec17, Sec17^FM>SS^, and Qc-SNAREs were expressed in *E. coli* and purified as described (Schwartz and Merz, 2009). Sec18 was expressed in *E. coli* and purified as described (Mayer et al., 1996). Full-length Vam3, Vti1 and Nyv1 were expressed in *E. coli* and purified as described (Zick et al., 2015; Mima et al., 2008; Zucchi and Zick, 2011). Ypt7 and HOPS were overproduced in yeast and purified as described (Zick and Wickner, 2013). Cy5-strepavidin was purchased from KPL, unlabeled avidin from Thermo Scientific, and R-phycoerythrin-biotin from Life Technologies. Monoclonal antibodies against ALP (Pho8), CPY (Prc1) and PGK1 were purchased from Molecular Probes (Life Technologies). Affinity-purified antibodies against Vam3, Vam7, Nyv1, Vti1, Sec17, Sec18, and Vps33 were as described (Schwartz and Merz, 2009). Lipids were purchased from Avanti Polar lipids, except for ergosterol (Sigma-Aldrich) and fluorescent lipids (Life Technologies).

### Microscopy

For fluorescent labeling of yeast vacuoles (Vida and Emr, 1995), cell cultures were shaken at 30°C and grown to early logarithmic phase (OD_600_ = 0.3 to 0.6). Cells were pelleted and resuspended in synthetic media supplemented with 5 μM FM4-64 (Life Technologies), then incubated 20 min (Fig 4, Supplemental Figs) or 3 hours (Fig 3) at 30°C. Labeled cells were rinsed once in synthetic media before resuspension in synthetic complete or dropout media and shaking at 30° C for 30 to 60 min. Cells were maintained in logarithmic phase (OD_600_ = 0.5 to 0.8) until observation by microscopy. Epifluorescence microscopy was performed as described (Paulsel et al., 2013).

### Vacuole protein sorting analysis

10 mLs of cells were grown to OD_600_ = 1.0 in synthetic complete media lacking appropriate nutrients to maintain plasmid selection. Cells were retrieved by centrifugation, suspended in 100 μL 1× SDS loading buffer with 100 μL glass beads, heated to 95° C for 10 min, and vortexed for 5 min to disrupt the cell wall and cell membrane. Cell extracts were separated from glass beads by centrifugation at 1000×g, and then insoluble material was removed by centrifugation at 20,000×g. Samples were analyzed by SDS-PAGE and western blotting.

### Immunoprecipitation

Affinity-purified Vam3 antibodies were covalently coupled to Protein A agarose beads using dimethyl pimelimidate (DMP; Pierce) as described (Harlow and Lane, 1999). Logarithmic phase cultures were harvested and spheroplasted. Briefly, 20 OD_600_-mL of cells were sedimented then resuspended in 2 mL 0.1 M Tris·Cl pH 9.4, 10 mM DTT for 10 min at room temperature. Cells were sedimented and resuspended in 4 mL spheroplasting buffer (yeast nitrogen base, 2% (w/v) glucose, 0.05% (w/v) casamino acids, 1M sorbitol, 50 mM Na·HEPES pH 7.4), then incubated with lyticase (re-purified Zymolyase 20T; Sekigaku) at 30°C for 30 min. Spheroplasts were sedimented once in spheroplasting buffer and resuspended in 2 mL ice cold lysis buffer (20 mM Na·HEPES pH 7.4, 100 mM NaCl, 20% (w/v) glycerol, 2 mM EDTA, 1 μg/mL aprotinin, 1 μg/mL leupeptin, 1 μg/mL pepstatin, 0.1 mM Pefabloc-SC, 1 mM PMSF, and a protease inhibitor cocktail (Roche), and lysed by ∼ 30 strokes with an ice-cold dounce homogenizer. Cell lysates were supplemented with 1% (v/v) Anapoe X-100 (Anatrace) and nutated at 4° C for 15 min. Insoluble debris was removed by centrifugation at 20,000×g for 15 min at 4° C. Clarified lysate was mixed with anti-Vam3-protein A beads and nutated for 30 min at 4° C. Beads were recovered by low speed centrifugation and rinsed 4 times in lysis buffer containing 0.5% (v/v) Anapoe X-100. Bound proteins were eluted by boiling in SDS-PAGE sample buffer (Laemmli, 1970). Unbound proteins remaining in the cell lysates were precipitated by addition of 1/10 vol. 0.15% deoxycholate and 1/10 vol. 100% vol. TCA, rinsed twice in acetone and resuspended in sample buffer (40 μL per OD_600_^×^mL equivalent). Samples were separated by SDS-PAGE, electroblotted to nitrocellulose, probed with primary antibodies as indicated in the figures and secondary antibodies as recommended by the manufacturer (LiCor), and analyzed on a LiCor Odyssey imaging system.

### In vitro fusion assays

Vacuole fusion assays were performed exactly as described (Schwartz and Merz, 2009). Dose-activity curves were fit with the Hill single-site model with adjustable slope, by the nonlinear least-squares method (GraphPad Prism). RPLs were formed by dialysis from β-octylglucoside proteolipid mixed micelles as in (Zick et al., 2015), in the configurations shown in Fig. S2B. A defined lipid composition highly similar to that of the yeast vacuole was employed (vacuolar membrane lipids, (VML; (Zick et al., 2014). The protein:lipid ratios were: 1:1000 for Qa, Qb, and R SNAREs, 1:2000 for Ypt7. RPL fusion assays were set up as described (Zick et al., 2015), but bovine serum albumin was omitted from the reactions, except for the reactions shown in Fig. 5E and F, where it was present. Reaction kinetics were fit with a second-order association model by the nonlinear least squares method using GraphPad Prism.

## Acknowledgements

We thank members of our groups for insightful discussions and comments on the manuscript, W. Wickner for hosting A.J.M. to learn the liposome fusion system and for provision of reagents, and R. Plemel for outstanding technical support. The Merz lab’s work on membrane fusion is supported by NIH-NIGMS GM077349; the Wickner lab’s, by GM023377.

---

**Fig. S1.**
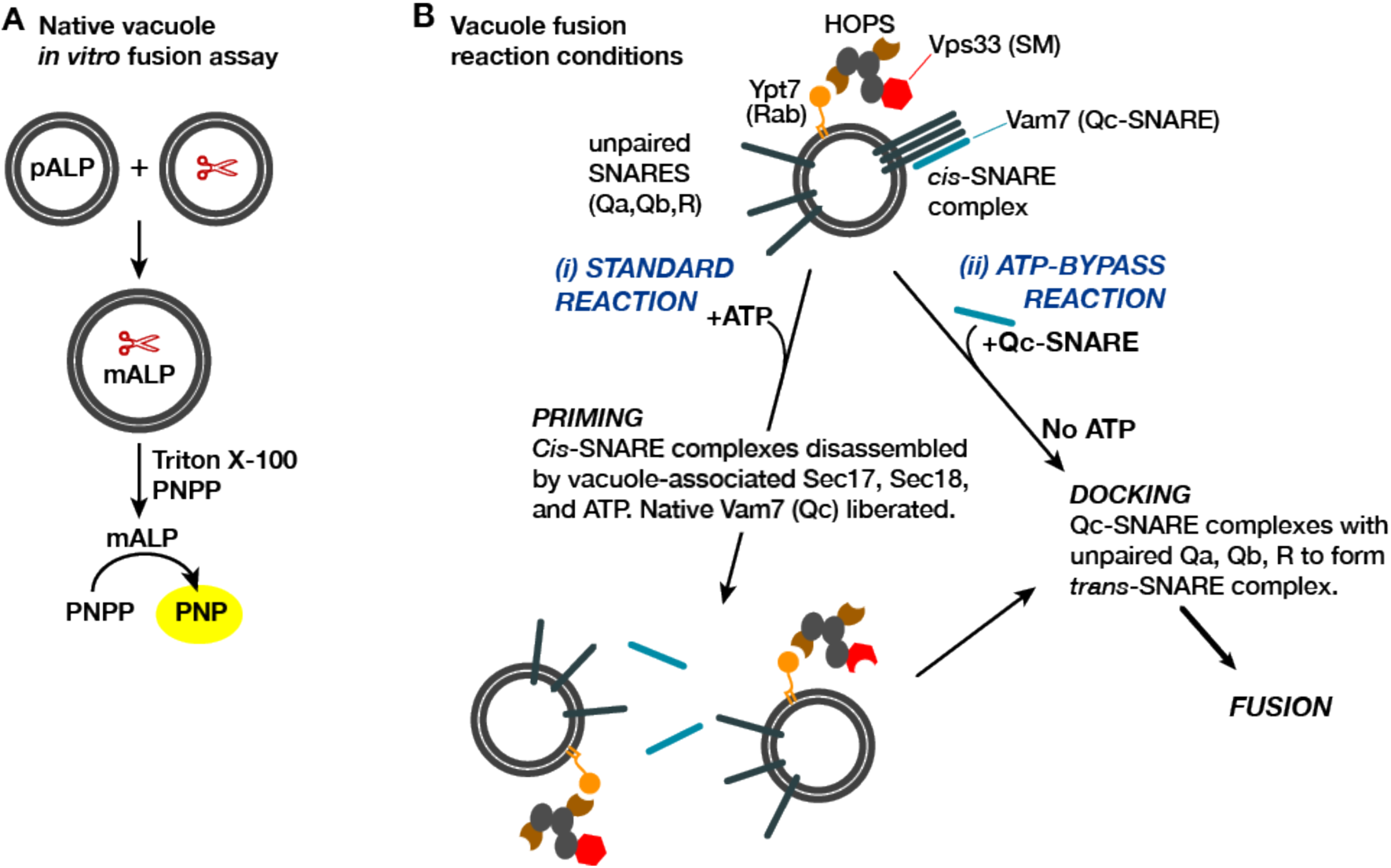
Vacuole fusion assay and reaction schemes used in this study. **A.** Content mixing assay system. One set of vacuoles is isolated from a strain containing the inactive vacuolar hydrolase proALP. The second set of vacuoles is isolated from a strain containing the maturases PrA and PrB (denoted by scissors). Content mixing causes proteolysis of proALP to mature mALP. The amount of mALP is assayed by lysis of the vacuole and addition of the colorimetric substrate para-nitrophenolphosphate, which is hydrolyzed to a yellow product by mALP. The formation of 4-nitrophenolate (PNP) is measured by spectrophotometry. **B**. Fusion assay configurations used in this study. The isolated vacuole bears active Ypt7-GTP, the SM-tether complex HOPS, and small amounts of both Sec17 and Sec18. SNAREs are present in two forms: unpaired SNAREs, and *cis*-SNARE complexes. In purified vacuole preparations, the Qc Vam7 is present only in the *cis*-SNARE complexes (Boeddinghaus et al., 2002). Thus, there are two ways to drive fusion of isolated vacuoles. In the standard “+ATP” reaction (i), ATP and Sec18 liberate SNAREs including native Vam7 from *cis*-complexes, leading to docking and fusion (Boeddinghaus et al., 2002; Mayer et al., 1996). In the “ATP bypass” reaction (ii), purified Vam7 (or Qc-Δ) is supplied in the absence of ATP, allowing docking and fusion to proceed.

**Fig. S2.**
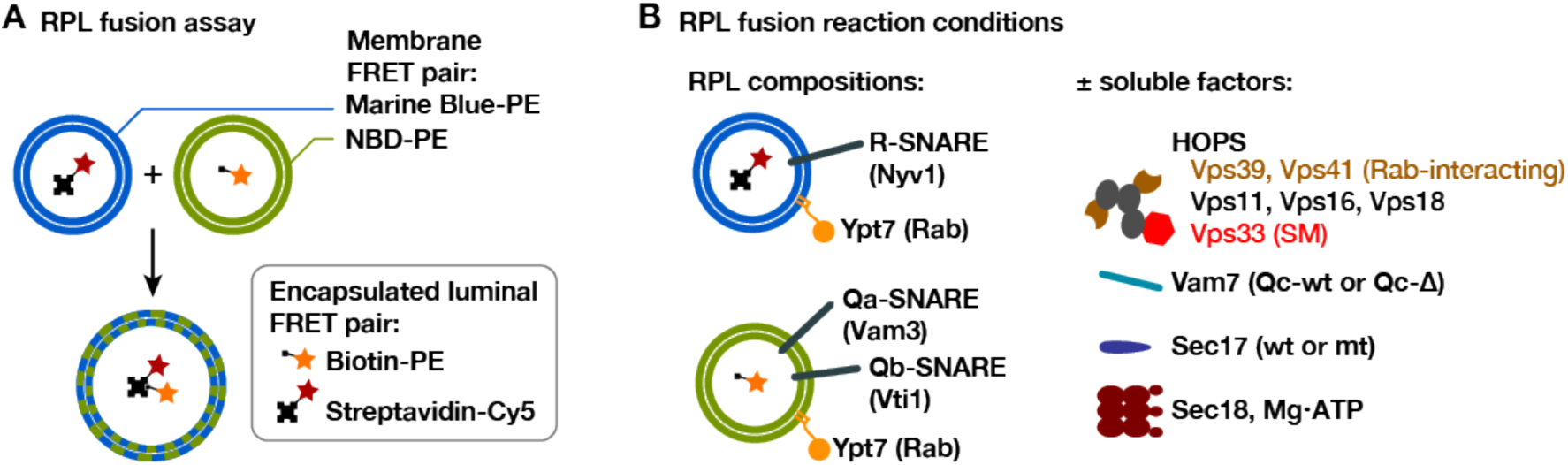
RPL fusion assay system. The RPL fusion assay simultaneously monitors lipid and content mixing using orthogonal FRET pairs (Zucchi and Zick, 2011). Lipid mixing is monitored by FRET between phosphatidylethanolamine (PE) lipid derivitized with either NBD or Marine Blue. Content mixing is monitored through the association of encapsulated biotin and avidin conjugates (to phycoerythrin and Cy5 fluorophores, respectively). Although both lipid and content mixing signals were collected in the experiments shown, only content mixing signals are presented because the two signals were highly correlated and because content mixing is the reaction’s biologically relevant endpoint. **B**. The RPLs used in this study contained a lipid mixture approximating that of the yeast vacuole membrane (Zick et al., 2014), native purified Ypt7-GTP, and either Qa- and Qb-, or R-SNAREs. The asymmetric SNARE topology allowed docking and fusion without any prior requirement for Sec18-mediated *cis-* SNARE complex disassembly.

**Fig. S3.**
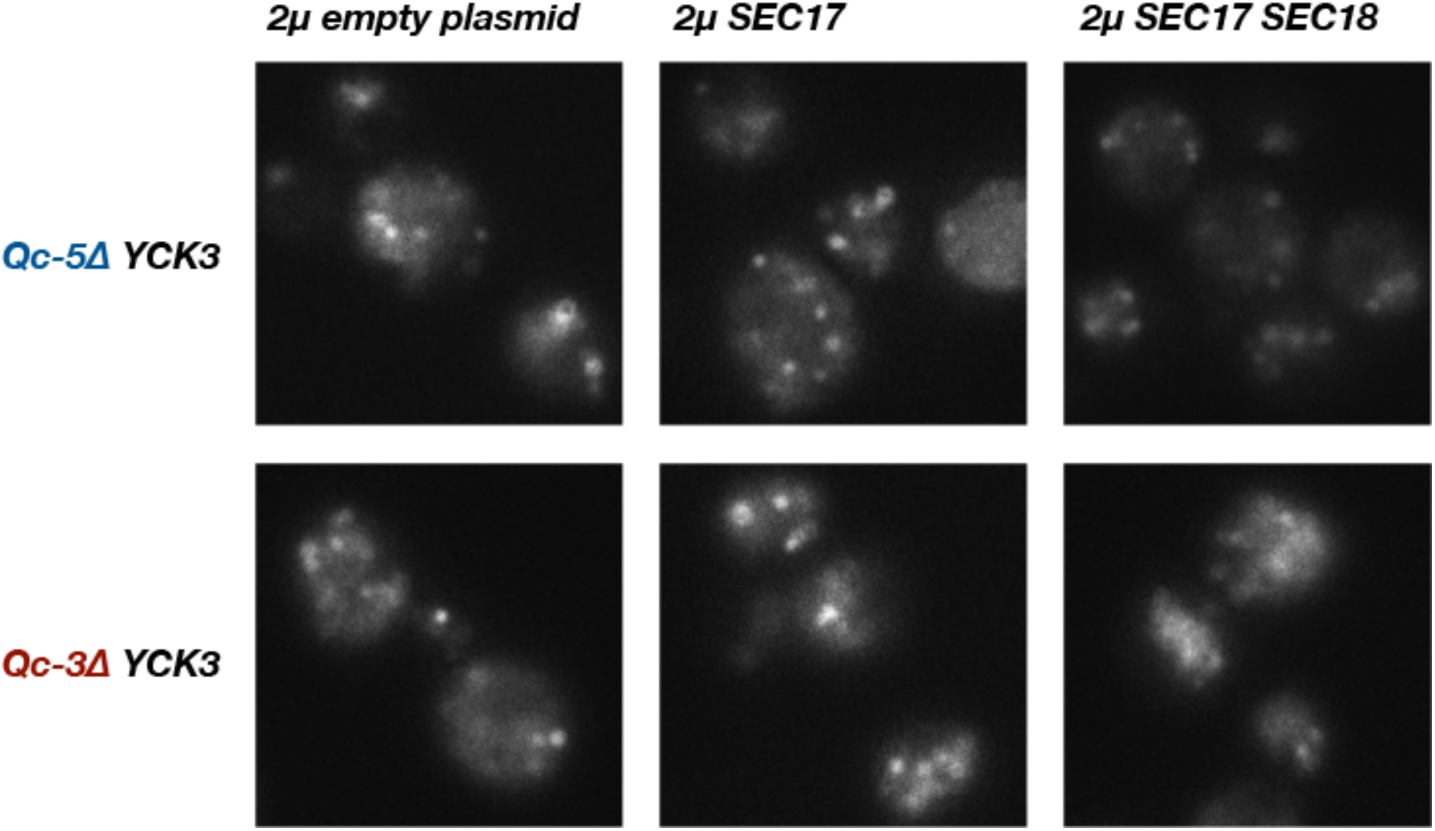
Little or no rescue of fragmented vacuole morphology was observed with Qc-3Δ or Qc-5Δ chromosomal integrants when Sec17 or Sec17 and Sec18 were overproduced in a ***YCK3*** genetic background. Vacuole morphology was assessed by FM4-64 staining as in Fig. 2. Representative fields are shown.

**Fig. S4.**
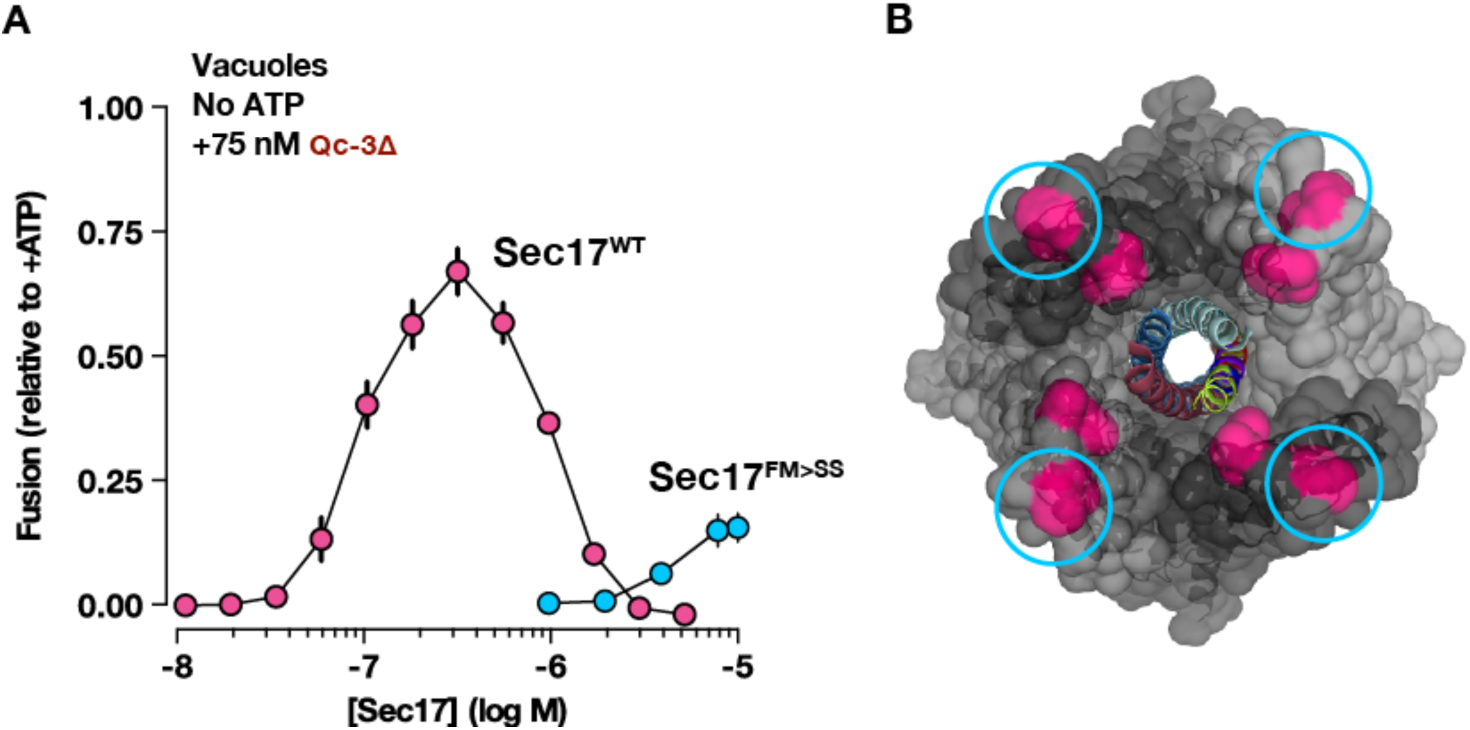
Membrane penetration facilitates Sec17 rescue of QcΔ-mediated homotypic vacuole fusion ***in vitro*.** **A**. Vacuole fusion reactions *in vitro* were assembled in the gain-of-function “ATP-bypass” configuration, using 75 nM Qc-3Δ, and the indicated concentrations of either Sec17 or Sec17^FM>SS^. Each point shows the mean ± s.e.m. of 3 independent experiments.

**Supplementary Table ST2.**
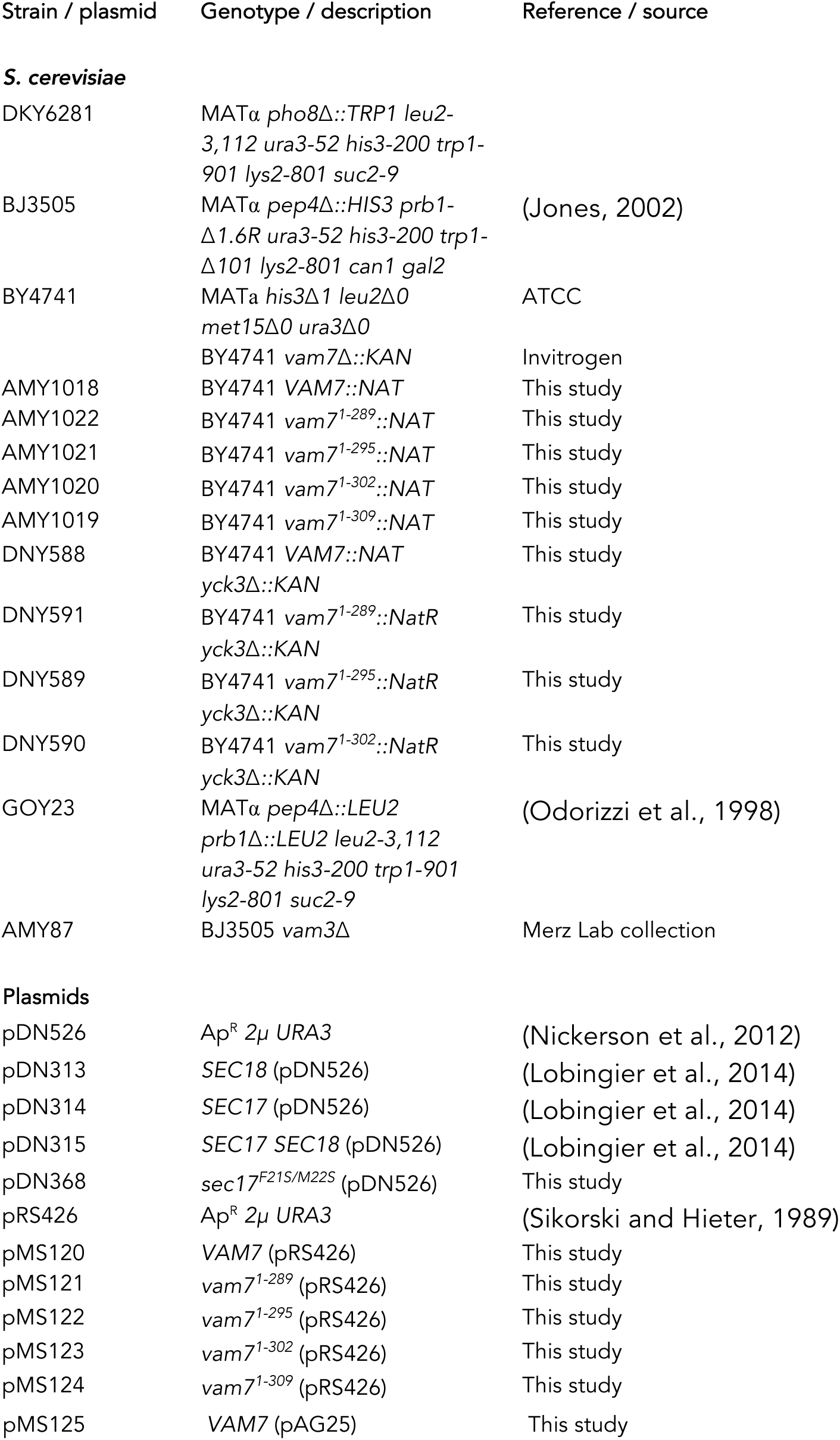
Strains and plasmids used in this study

